# Crosstalk between AML and stromal cells triggers acetate secretion through the metabolic rewiring of stromal cells

**DOI:** 10.1101/2021.01.21.427406

**Authors:** Nuria Vilaplana-Lopera, Ruba Almaghrabi, Grigorios Papatzikas, Vincent Cuminetti, Mark Jeeves, Elena González, Alan Cunningham, Ayşegül Erdem, Frank Schnuetgen, Manoj Raghavan, Sandeep Potluri, Jean-Baptiste Cazier, Jan Jacob Schuringa, Michelle AC Reed, Lorena Arranz, Ulrich L Günther, Paloma Garcia

## Abstract

Acute myeloid leukaemia (AML) cells interact and modulate components of their surrounding microenvironment into their own benefit. Stromal cells have been shown to support AML survival and progression through various mechanisms. Nonetheless, it is unclear whether AML cells could establish beneficial metabolic interactions with stromal cells. Here, we identify a novel metabolic crosstalk between AML and stromal cells where AML cells prompt stromal cells to secrete acetate for their own consumption to feed the tricarboxylic acid cycle (TCA). By performing transcriptome analysis, tracer-based metabolic NMR analysis and ROS measurements, we observe that stromal cells present a higher rate of glycolysis. We also find that acetate in stromal cells is derived from pyruvate via chemical conversion under the influence of reactive oxygen species (ROS) following ROS transfer from AML to stromal cells via gap junctions. Overall, we present a unique metabolic communication between AML and stromal cells that could potentially be exploited for adjuvant therapy.

## Introduction

Acute myeloid leukaemia (AML) is a heterogeneous clonal disease characterised by a rapid proliferation of aberrant immature myeloid cells that accumulate in the bone marrow, and eventually in the blood and other organs, severely impairing normal haematopoiesis. AML cells show a highly adaptive metabolism that allows them to efficiently use a variety of nutrients to obtain energy and generate biomass (reviewed in ^1^). This high metabolic plasticity confers AML cells a strong advantage against normal haematopoietic cells and has been related to AML aggressiveness. Although the metabolism in AML cells has been broadly investigated ^1^, fewer studies have focused on identifying metabolic alterations related to the interaction between AML and niche cells. For instance, AML cells are known to interact and modulate niche components for their own support by secreting soluble factors ^2–6^, via exosomes ^7–9^ or by establishing direct interactions, mediated by gap junctions ^10^ or tunnelling nanotubes ^11^. These interactions with components of the niche provide AML cells with survival cues, chemoresistance and increased relapse in AML patients ^2, 4, 7, 10, 12–14^. Furthermore, it has been reported that adipocytes in the niche secrete fatty acids, which are metabolised by AML cells through β-oxidation to obtain energy, protecting AML cells from apoptosis and ROS ^15, 16^.

As a consequence of their highly proliferative demand, leukaemic stem cells (LSCs) ^17–20^ and, particularly, chemotherapy-resistant AML cells ^21^ present abnormally high levels of reactive oxygen species (ROS) ^22, 23^. How AML cells cope with high ROS levels has been intriguing and recent reports are shedding some light on whether the microenvironment plays a role in the redox metabolism of AML cells. For instance, it was reported that Nestin^+^ bone marrow mesenchymal stem cells (BMSCs) support AML progression by increasing the bioenergetic capacity of AML cells and providing them with glutathione (GSH)-mediated antioxidant defence to balance the excess ROS ^14^. Similarly, a recent study showed that co-culturing BMSCs with AML cells leads to a decrease in AML ROS levels due to an activation of the antioxidant enzyme GPx-3 in AML cells ^24^.

Our work provides new insight into the metabolic and redox crosstalk between AML and stromal cells, revealing a new metabolic interaction between AML and stromal cells. By combining transcriptomic and nuclear magnetic resonance (NMR) data, our results demonstrate that stromal cell metabolism is rewired in co-culture giving higher glycolysis and pyruvate decarboxylation, leading to acetate secretion. Our results also show that AML cells are able to transfer ROS to stromal cells by direct interaction through gap junctions and that these ROS can be used by stromal cells to generate and secrete acetate, which is utilised by AML cells as a biofuel by metabolising it via the TCA cycle. Targeting ROS transfer via modulation of gap junctions to suppress the antioxidant protection provided by stromal cells could serve as an adjuvant therapy to eradicate AML.

## Results

### Co-culturing AML and stromal cells in direct contact triggers acetate secretion by stromal cells

We first sought to determine whether interactions between AML and stromal cells in co-culture would result in differences in the consumption or production of extracellular metabolites. For this purpose, three AML cell lines (SKM-1, Kasumi-1 and HL-60) were cultured separately and in co-culture with MS-5, a stromal cell line capable of maintaining hematopoiesis ^25^, and the metabolic composition of the extracellular medium was analysed by ^1^H-NMR and compared at different time points. The most striking difference found in co-culture compared to cells cultured separately was an increased secretion of acetate, which was common for the three cell lines used (Fig. 1a,b). Moreover, only stromal cells secreted acetate to a lower extent when cultured alone whereas AML cells did not secrete any acetate when cultured alone. Altogether, these findings suggest that acetate secretion is indeed a result of a direct interaction between AML and stromal cells. In addition, the observation that only stromal cells secrete acetate under these conditions suggests that stromal cells could be responsible for the increased acetate secretion found in co-culture.

**Fig 1.**
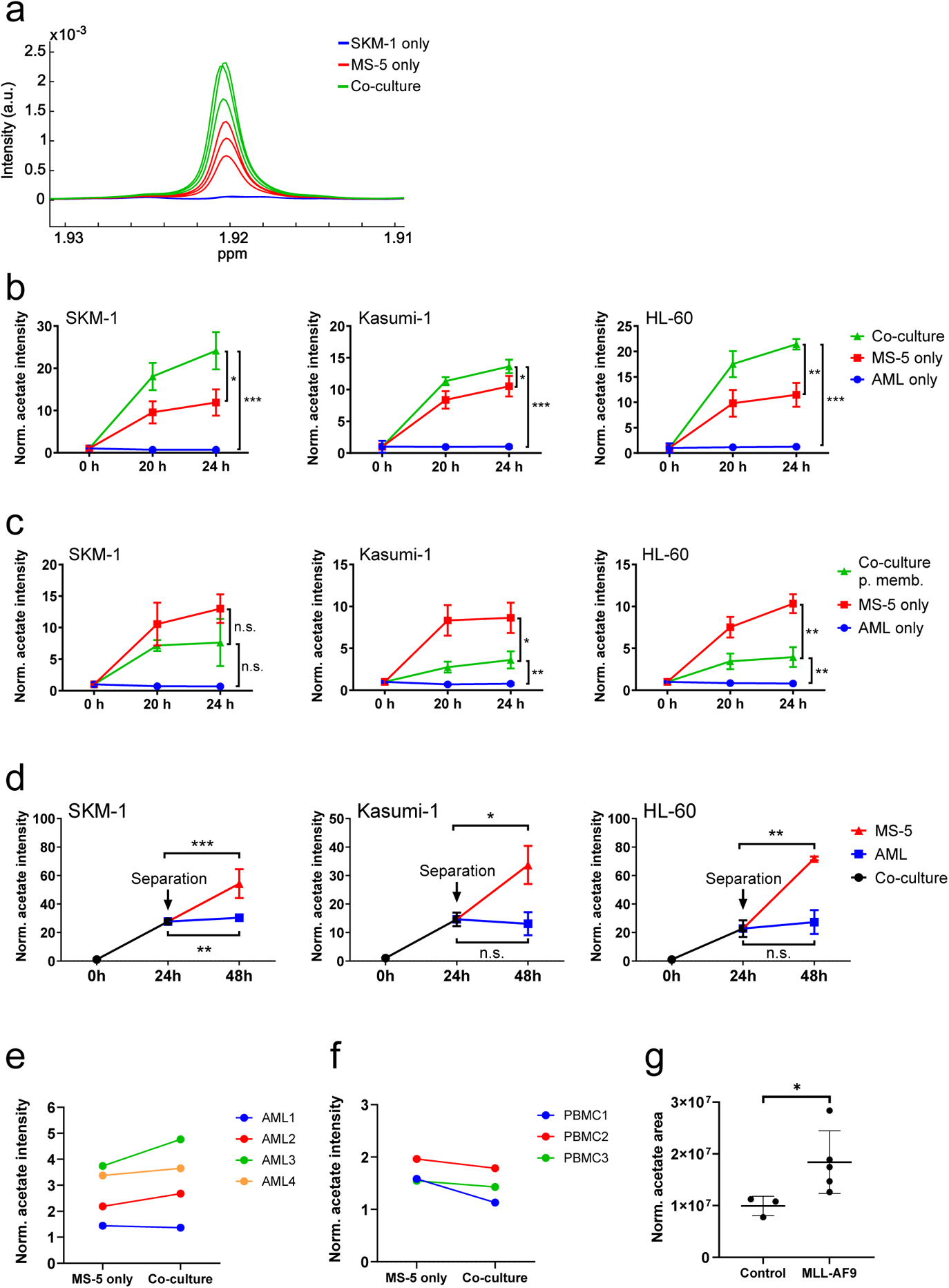
Acetate secretion by stromal cells increases in AML-stroma co-cultures of several AML cell lines and primary AML cells in direct contact. **a**, Section of ^1^H-NMR spectra, corresponding to the methyl group of acetate, from extracellular medium samples of SKM-1 cells cultured alone (blue), MS-5 cells cultured alone (red) and SKM-1 and MS-5 cells in co-culture (green) after 24 hours. **b**, Extracellular acetate levels in AML cell lines (SKM-1, Kasumi-1 and HL-60) cultured alone (blue), MS-5 cells cultured alone (red) and AML and MS-5 cells in co-culture in direct contact (green) at 0, 20 and 24 hours of incubation. Each point represents the mean of n=3 independent experiments and error bars represent standard deviations. **c**, Extracellular acetate levels in AML cell lines cultured alone (blue), MS-5 cells cultured alone (red) and AML and MS-5 cells in co-culture separated by a 0.4 µm permeable membrane (green) at 0, 20 and 24 hours of incubation. Each point represents the mean of n=3 independent experiments and error bars represent standard deviations. **d**, Extracellular acetate levels in AML cell lines and MS-5 cells in co-culture (black) for 24 hours and after being separated and cultured alone in the same medium until 48 hours (blue for AML and red for MS-5). Each point represents the mean of n=3 independent experiments and error bars represent standard deviations. For **b**, **c, d** and **g** unpaired Student’s t-tests were applied for each condition (black brackets) and p-values are represented by n.s. for not significant * for p-value<0.05, ** for p-value<0.01 and *** for p-value<0.001. **e**, Extracellular acetate levels in MS-5 cells cultured alone (black) and primary patient-derived AML cells co-cultured with MS-5 cells (grey) at 48h. Each set of points represents an independent experiment (n=4). **f**, Extracellular acetate levels in MS-5 cells cultured alone (black) and healthy donor-derived peripheral blood mononuclear CD34^+^ (PBMC) cells co-cultured with MS-5 cells (grey) at 48h. Each set of points represents an independent experiment (n=3). **g**, Acetate levels in bone marrow extracellular fluid of C57BL6/J mice 6 months after transplantation with bone marrow nucleated cells isolated from control or MLL-AF9 transgenic mice. For b, c, d and g unpaired Student’s t-tests were applied for each condition (black brackets) and p-values are represented by n.s. for not significant * for p-value<0.05, ** for p-value<0.01 and *** for p-value<0.001.

Moreover, we examined the levels of other common extracellular metabolites, including glucose, lactate, glutamate and glutamine. As shown in Supplementary Fig. 1a higher consumption of glucose along with a higher secretion of lactate was observed in AML and stromal cells in co-culture compared to single cultures, suggesting a higher glycolytic flux in co-culture. However, the levels of glucose consumption and lactate production in co-culture were similar to the sum of the glucose consumption and lactate production levels of the AML and stromal cells in single cultures, suggesting that the overall increase in glycolysis in co-culture was just a result of culturing both cell types together. Additionally, we observed no variation in glutamate and glutamine levels suggesting that these metabolites are not involved in interactions that result from co-culture and are utilised depending on their availability.

The metabolic differences found in co-culture, could be due to altered proliferation under these conditions. To determine whether this was the case, a CFSE-based proliferation assay was performed in which AML cells and not stromal cells were stained and their growth was compared when cultured separately vs in co-culture. Over 48 hours, none of the AML cell lines tested presented differences in their proliferation rates (Supplementary Fig. 2a), thus confirming that the metabolic changes found in co-culture were caused by a mechanism independent from changes in proliferation.

**Fig. 2.**
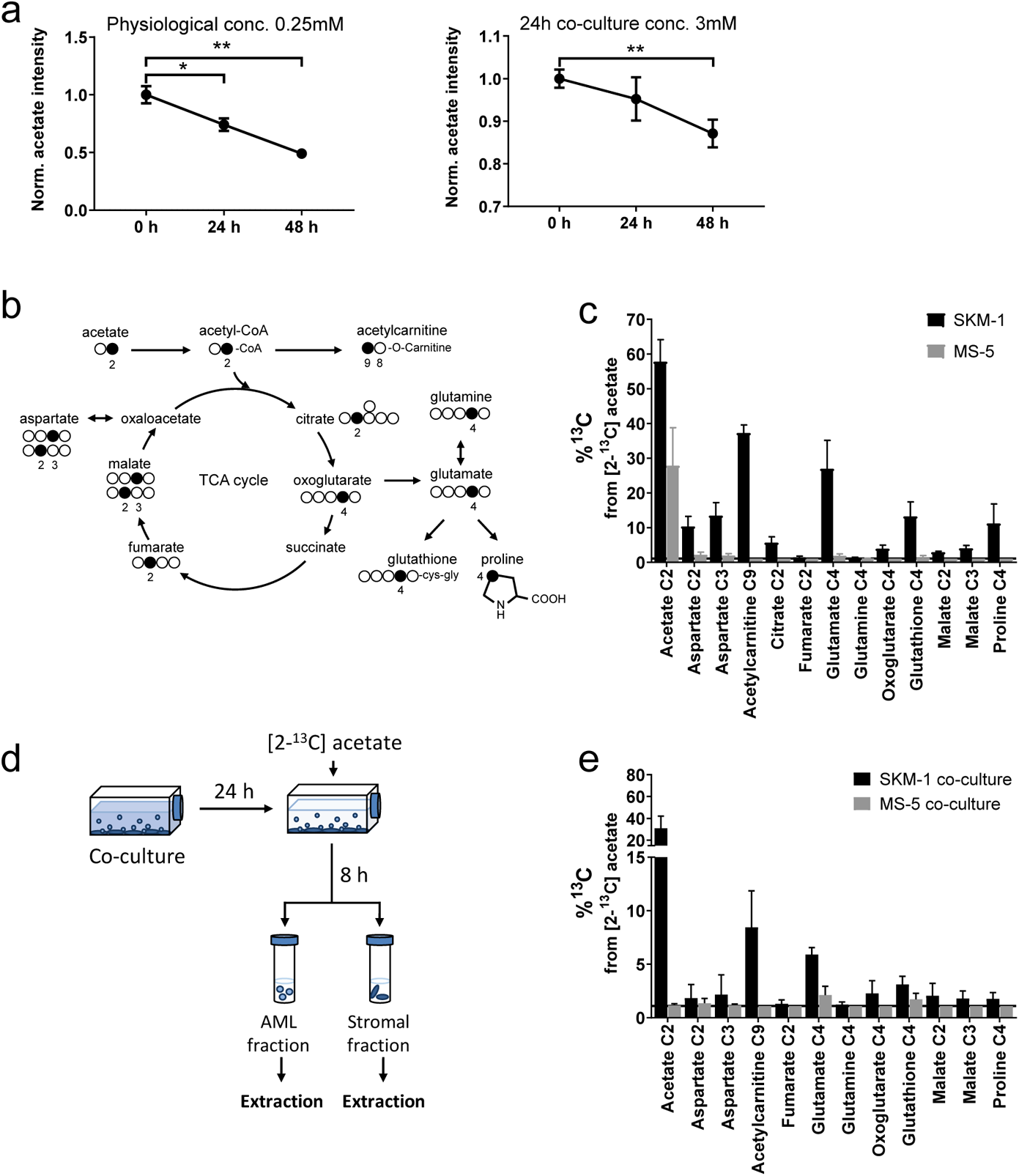
AML cells import the secreted acetate in co-culture to use it as a biofuel. **a,** Extracellular acetate levels in SKM-1 cells cultured with 0.25 mM and 3 mM acetate medium after 0, 24 and 48 hours. Each point represents the mean of n=3 independent experiments and error bars represent standard deviation. An unpaired Student’s t-test was applied for each condition (black brackets) and p-values are represented by * for p-value<0.05, ** for p-value<0.01 and *** for p-value<0.001. **b,** Scheme of label distribution arising from [2-^13^C]acetate in TCA cycle intermediates. Black circles correspond to expected labelled positions. **c,** ^13^C percentages of label incorporation in polar metabolites from labelled acetate in SKM-1 and MS-5 cells after two hours of incubation with [2-^13^C]acetate (4 mM) derived from ^1^H-^13^C-HSQC NMR spectra. Bars represent the mean of the ^13^C percentage and error bars represent the standard deviations for n=3 independent experiments. **d,** Experimental design for acetate labelling in co-culture. Briefly, AML cells are co-cultured with stromal cells for 24 hours prior to addition of [2-^13^C]acetate (extra 4 mM). After 8 hours of the addition of acetate, cells are separated, and metabolites are extracted. **e,** ^13^C percentages on polar metabolites in SKM-1 and MS-5 cells in co-culture. Cells were co-cultured for 24 hours before the addition of extra 4 mM sodium [2-^13^C]acetate. Bars represent the mean of the ^13^C percentage and error bars represent the standard deviations of n=3 independent experiments. For C and E, ^13^C natural abundance is represented as a black bar at %^13^C = 1.1.

Next, we aimed to determine whether cell-to-cell contact could play a role in the increased acetate secretion found in co-culture. For this we co-cultured AML and stromal cells separated by a permeable membrane, allowing cells to share the extracellular medium but impeding cell-to-cell contact. Co-culturing cells using a permeable membrane blocked the increase in acetate secretion observed under direct contact conditions (Fig. 1c). In fact, cells in co-culture presented lower levels of acetate than MS-5 cells cultured alone revealing that direct cell-to-cell contact is required for acetate secretion in co-culture.

To examine whether increased acetate secretion is specific for the interaction of AML cells with stromal cells, we co-cultured AML cells with the cervical cancer cell line HeLa and compared the levels of acetate of each cell type cultured alone. We found that there was no acetate secretion in co-culture, also not by HeLa and AML cells cultured alone (Supplementary Fig. 2b), suggesting that increased acetate secretion could be specific for the AML-stromal interaction.

Considering that our previous data seemed to indicate that MS-5 cells were responsible for acetate secretion when co-cultured with AML cell lines, we decided to investigate how the levels of extracellular acetate would vary after separating cells from co-culture. The three AML cell lines were co-cultured for 24h with MS-5 cells prior to separation and were subsequently cultured in the same spent media. Extracellular acetate levels in previously co-cultured MS-5 cells followed a similar trend as before separation, suggesting that it is most likely that MS-5 cells are responsible for the increased acetate secretion found in co-culture (Fig. 1d). Moreover, AML cells did not follow this trend as they either maintained the levels of acetate seen prior to separating the cells (Kasumi-1 and HL-60) or presented only a moderate increase (SKM-1) after being separated from co-culture.

We further investigated whether increased acetate secretion could take place in primary co-cultures using AML cells derived from patients and whether acetate secretion in co-culture could be specific for AML cells. To address this question, we isolated the CD34^+^ population of four primary AML patient samples, and three independent healthy donors, cultured them alone vs in co-culture with MS-5 cells, and analysed the composition of the extracellular medium at different time points. We found that three out of four primary AML samples presented higher levels of acetate when co-cultured with MS-5 cells compared to both MS-5 and AML cells cultured alone (Fig. 1e). Contrary, none of the healthy donor samples showed an increased acetate secretion in co-culture suggesting that acetate secretion is specific for AML cells in co-culture (Fig. 1f).

Moreover, we sought to determine whether in an *in vivo* setting, increased acetate production would be observed. For this, acetate levels were analysed in the bone marrow extracellular fluid (BMEF) of mice transplanted either with MLL-AF9 leukaemic cells or with healthy wild type hematopoietic cells. These experiments revealed that a significantly higher amounts of acetate was present in the BMEF of mice suffering from leukaemia compared to controls (Fig. 1g).

### AML cells consume and use acetate secreted by stromal cells to feed the TCA cycle

Following the finding that stromal cells are responsible for acetate secretion in co-culture, we decided to determine whether AML cells might be able to metabolise the secreted acetate. We first sought to define the concentration of secreted acetate in the extracellular medium in co-culture. For this, we compared a sample of extracellular medium from a co-culture of SKM-1 and MS-5 cells after 24 hours to a calibration curve (Supplementary Fig. 2c) which allowed us to determine that the concentration of acetate in co-culture, which was approximately of 3 mM. We then investigated whether SKM-1 cells can consume acetate both in physiological and co-culture concentrations. In both cases SKM-1 cells consumed acetate after 48 hours, and in physiological conditions also after 24 hours (Fig. 2a).

We used a tracer-based approach using [2-^13^C]acetate to assess whether SKM-1 or MS-5 cells can utilise acetate (Fig. 2b,c and Supplementary Fig. 3a). Both cell types import ^13^C labelled acetate as observed in NMR spectra (Fig. 2c). However, only SKM-1 cells showed ^13^C label incorporation in several TCA cycle related metabolites, including aspartate, citrate, glutamate, 2-oxoglutarate, glutathione and proline, as well as in acetylcarnitine. Acetylcarnitine is known to be produced by cells when large amounts of acetyl-CoA are present in the mitochondria ^26, 27^, which could be in line with this experiment in which a 4 mM concentration of [2-^13^C]acetate was used. Overall, this data suggests that acetate in co-culture could be utilised by SKM-1 cells but not by MS-5 cells.

**Fig. 3.**
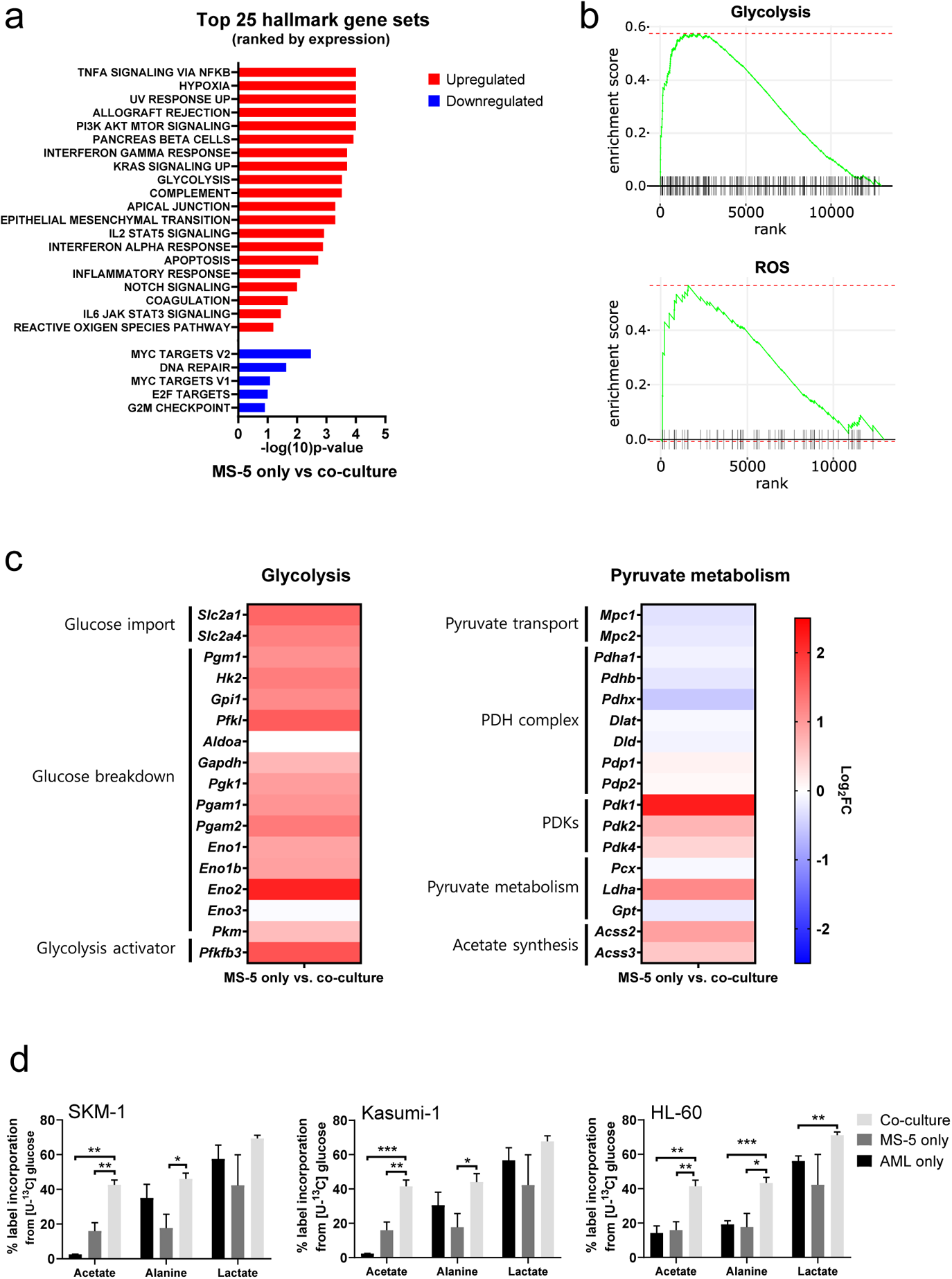
Transcriptomic data reveals that stromal cell metabolism is shifted towards higher glycolysis and ROS upon co-culture with AML cells. **a,** Top 25 GSEA hallmark gene sets ranked by expression in MS-5 cells only vs co-culture with SKM-1 cells, analysed using the collection of hallmark gene sets from Molecular Signature Database with a false discovery rate threshold at 5%. P-values for each pathway are presented as –log(10)p-value. **b,** GSEA enrichment score plots of glycolysis and ROS generated using Sleuth 0.30.0 R statistical package. **c,** Fold change values of detected gene transcripts (TPMs) related to glycolysis and pyruvate metabolism. FC values are represented as log_2_FC, red values indicate upregulation and blue values indicate downregulation in MS-5 cells in co-culture. **d,** Label incorporation from [U-^13^C]glucose into extracellular metabolites in AML and MS-5 cells cultured alone or in co-culture after 24 hours. Bars represent the mean of n=3 independent experiments and error bars represent standard deviations.

In order to determine whether AML cells can import and metabolise the secreted acetate in co-culture, we co-cultured AML and MS-5 cells for 24 hours, we then added [2-^13^C]acetate to the spent extracellular medium and cultured cells for 8 hours before analysing the intracellular metabolites in each cell type (Fig. 2d). We found that only AML cells presented ^13^C labelling in intracellular metabolites (Fig. 2e, Supplementary Fig. 3b). Moreover, the metabolisation pattern for all AML cells was similar to that observed for SKM-1 cells alone (Fig. 2c), where label was incorporated into TCA cycle-related metabolites. Overall, these results revealed that AML cells uptake and utilise the acetate secreted by stromal cells in co-culture as a substrate to feed into the TCA cycle.

### Transcriptomic data highlights a metabolic rewiring of stromal cells in co-culture characterised by upregulation of glycolysis and downregulation of pyruvate dehydrogenase

After establishing that stromal cells are responsible for acetate secretion (Fig. 1d) and that AML cells consumed the secreted acetate in co-culture (Fig. 2d), we sought to elucidate the mechanism behind acetate secretion by MS-5 cells in co-culture. Thus, we set out to perform global gene expression profiling by RNA-seq comparing cells cultured alone and in co-culture (Supplementary Fig. 4a). With this approach we identified 587 differentially expressed genes (q-value<0.1) (Supplementary Fig. 4b); with 476 genes being upregulated and 111 genes being downregulated in co-culture.

**Fig. 4.**
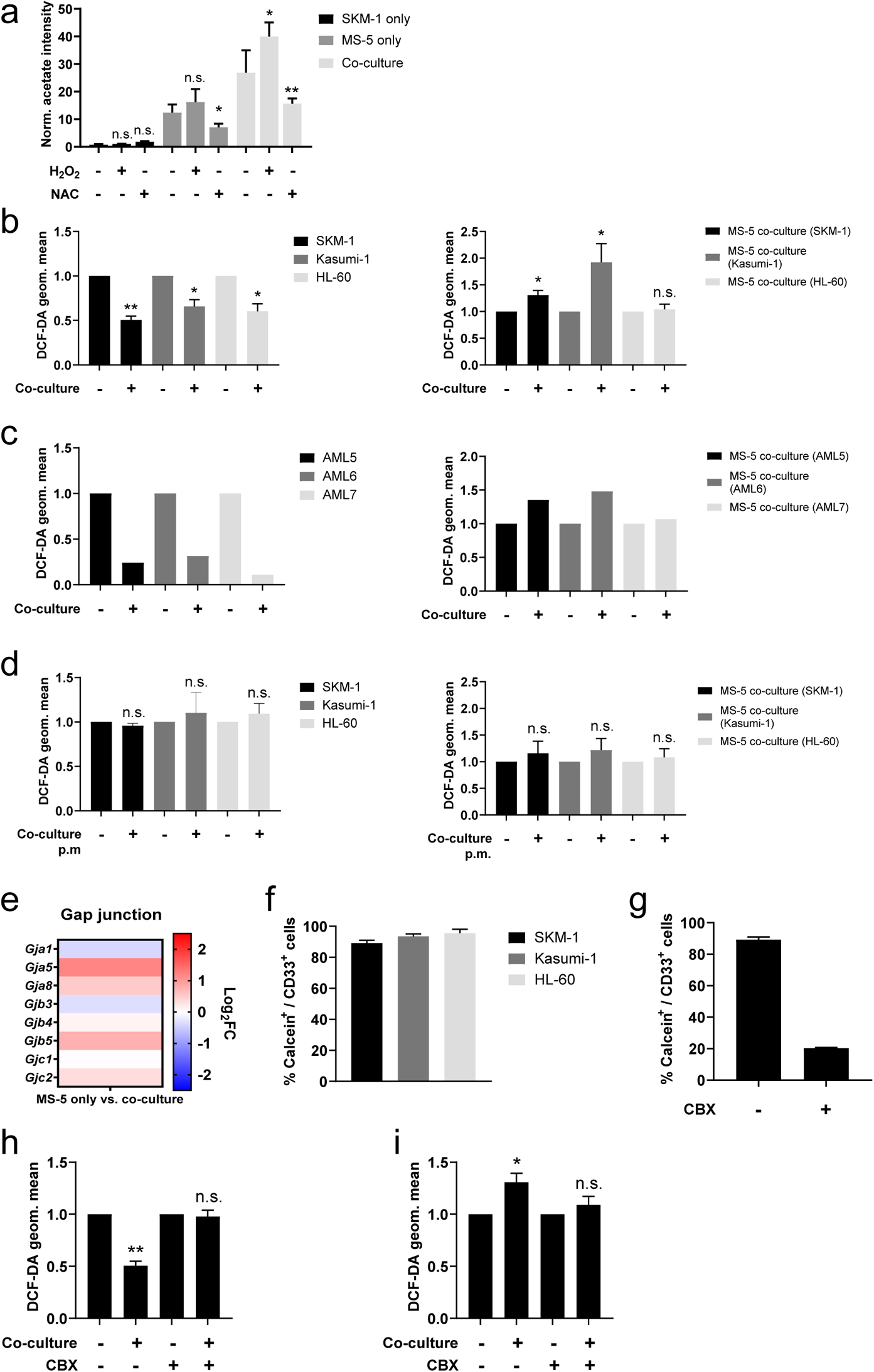
Acetate secretion is linked to ROS transfer from AML to stromal cells via gap junctions. **a,** Extracellular acetate levels in SKM-1 (black) and MS-5 cells cultured alone (dark grey) and in co-culture (light grey) for 24 hours in a control medium, medium with 50 µM H_2_O_2_ or medium with 5 mM NAC. **b**, **c** and **d** Intracellular ROS levels measured by H_2_DCFDA staining in **b** and **d** AML cells or **c** primary patient-derived AML cells and MS-5 cells cultured alone and in co-culture in **b** and **c** direct contact or **e,** fold change values of detected gene transcripts (TPMs) related to gap junctions. FC values are represented as log_2_FC, red values indicate upregulation and blue values indicate downregulation in MS-5 cells in co-culture. **f,** Frequencies of Calcein-AM and CD33 positive AML cell lines after being in co-culture with Calcein-AM stained MS-5 cells for 3 hours. **g,** Frequencies of Calcein-AM and CD33 positive SKM-1 cells (treated or untreated with 200 µM carbenoxolone for 24 hours) after being in co-culture with Calcein-AM stained MS-5 cells for 3 hours. **h** and **i** Intracellular ROS levels measured by H_2_DCFDA staining in **h** AML cells and **i** MS-5 cells cultured alone and in co-culture in direct contact treated or untreated with 200 µM carbenoxolone for 24 hours. **j,** Cell viability was assessed in SKM-1 cells cultured with 5-azacytidine or DMSO, with and without NAC over 48 hours by trypan blue staining and counting. Each point represents the mean and error bars represent the standard deviations of n=3 independent experiments. For **a**, **b**, **c**, **d**, **f**, **g**, **h** and **i** bars represent the mean of n=3 independent experiments and error bars represent standard deviations. For **a**, **b**, **d**, **h** and **i** and **j** unpaired Student’s t-test was applied for **a** each condition (H_2_O_2_/NAC) was compared to untreated; for **b**, **d**, **h** and **i** co-culture conditions were compared to cells cultured alone; for **j** each pair of conditions was compared (black brackets). P-values are represented by n.s. for not significant, * for p-value<0.05, ** for p-value<0.01 and *** for p-value<0.001.

Following clustering of differentially expressed genes, gene set enrichment analysis (GSEA) was performed (Fig. 3a) revealing a positive correlation with the expression of genes that are part of the glycolysis pathway (MSigDB: M5937) as well as the reactive oxygen species pathway (MsigDB: M5938) (Fig. 3b).

A closer examination of the genes involved in the glycolysis pathway revealed a major upregulation of several glycolysis-related genes in MS-5 cells in co-culture (Fig. 3c). Most of the genes involved in glucose transport (*Slc2a1* and *Slc2a4*) and glucose breakdown to pyruvate (*Pgm1*, *Hk2*, *Gpi1*, *Pfkl*, *Gapdh*, *Pgk1*, *Pgam1-2*, *Eno1-1b-2* and *Pkm*) were upregulated, including the gene encoding for the 6-phosphofructo-2-kinase/fructose-2,6-biphosphatase 3 (*Pfkfb3*), a well-known activator of glycolysis.

In contrast, individual examination of genes related to pyruvate metabolism revealed that the genes related to pyruvate conversion to acetyl-CoA were downregulated (Fig. 3c). Pyruvate transport into the mitochondria (*Mpc1-2*) and several pyruvate dehydrogenase (PDH) complex-related genes (*Pdha1*, *Pdhb* and *Pdhx*), involved in the conversion of pyruvate to acetyl-CoA, remained largely unaltered or were slightly downregulated by co-culture. Additionally, the pyruvate dehydrogenase kinases (*Pdk1*, *Pdk2* and *Pdk4*), which inhibit the activity of the PDH complex, were found to be upregulated by co-culture. Interestingly, the acetyl-CoA synthetases (ACSs), which can generate acetyl-CoA from acetate but have also been reported to perform the reverse reaction, encoded by *Acss2* and *Acss3*, were found to be slightly upregulated in co-culture. Pyruvate can also be converted to 2-oxaloacetate via pyruvate carboxylase (*Pcx*), to lactate by lactate dehydrogenase (*Ldha*), and to alanine via alanine transaminase (*Gpt*). Only *Ldha* was found to be moderately upregulated in co-culture. However, we did not observe a substantial increase in lactate production experimentally in co-culture. Altogether, transcriptomic data suggests that MS-5 cells in co-culture present a major upregulation of glycolysis and downregulation of the PDH complex.

To explore whether glycolysis is upregulated at a metabolic level in MS-5 cells in co-culture at the metabolic level and whether acetate could derive from glucose, we performed [U-^13^C]glucose tracing on AML and MS-5 cells cultured alone and in co-culture and analysed the label incorporation in glycolysis-related extracellular metabolites (Fig. 3d). For the three AML cell lines tested, extracellular acetate presented significantly higher label incorporations from [U-^13^C]glucose in co-culture compared to cell types cultured alone, providing evidence that the secreted acetate in co-culture derives from glucose. Lactate and alanine, which can be synthesised from pyruvate, did not show significant increases in label incorporation from [U-^13^C]glucose in co-culture compared to each cell type cultured alone for all the AML cell lines, with the exception of alanine in HL-60. Overall, these results are in agreement with the transcriptomic data (Fig. 3c), highlighting that glucose metabolism is upregulated in co-culture but also confirming that acetate derives from glycolysis.

### AML cells rewire stromal cell metabolism transferring ROS via gap junctions to obtain acetate

Tracer-based data on MS-5 cells in co-culture revealed that acetate secreted in co-culture derives from glucose (Fig. 3d). However, transcriptomic data on MS-5 cells in co-culture suggested that acetate did not derive from pyruvate via acetyl-CoA (Fig. 3c). Moreover, acetate secretion was also observed in MS5 cells grown in thiamine-free media (Supplementary Fig. 5a), confirming that the acetate secretion was not dependent on keto acid dehydrogenases ^28^ and indicating that a different mechanism must be responsible. An alternative mechanism of acetate synthesis involving a non-enzymatic oxidative decarboxylation of pyruvate into acetate had previously been described ^28–30^. This mechanism was reported to be mediated by ROS in mammalian cells and was linked to cells prone to overflow metabolism under the influence of high rates of glycolysis and excess pyruvate. Hence, we decided to investigate whether ROS might play a role in acetate secretion in our co-culture system.

We first modulated ROS levels in AML cells and MS-5 cells cultured alone and in co-culture and measured acetate production. Hydrogen peroxide was used to increase ROS levels and N-acetylcysteine (NAC) was used as a ROS scavenger. Extracellular acetate levels were measured by ^1^H-NMR after 24 hours. Increasing ROS levels with peroxide resulted in a significant increase in acetate secretion, particularly in SKM-1 and Kasumi cells in co-culture (Fig. 4a, Supplementary Fig. 5b)). This experiment could not be carried out with HL-60 cells as peroxide treatment severely impaired the viability of HL-60 cells, as previously reported ^31^. Additionally, when the ROS scavenger NAC was used, a significant decrease in the levels of acetate in both MS-5 cells cultured alone and in all the co-cultured cell lines was observed (Fig. 4a, Supplementary Fig. 5b). Interestingly, the production of acetate in cells in co-culture treated with NAC recovered the phenotype of MS-5 cells cultured alone, indicating that acetate synthesis in MS-5 cells in co-culture is mediated by ROS (Fig. 4a).

Next, we compared intracellular ROS levels in AML and stromal cells in co-culture and cultured alone by labelling cells with the H_2_DCFDA fluorescent dye. Fluorescence measurements showed that ROS levels in the three AML cell lines used were significantly decreased in co-culture, whilst in the stromal cells ROS levels were significantly increased in two of the three co-cultures (Fig. 4b). These results suggested that AML cells might transfer ROS to stromal cells. We also performed the same experiment using three primary AML samples to corroborate the previous result. Fluorescence analysis showed decreased ROS levels in AML samples in co-culture and increased ROS levels in MS-5 cells in co-culture for the three primary AML samples analysed (Fig. 4c), suggesting ROS transfer from AML cells to stromal cells in co-culture.

To further test whether AML cells might transfer ROS through a contact-dependent mechanism, we compared the intracellular ROS levels in AML and stromal cells cultured alone and co-cultured without direct contact using a permeable membrane. Fluorescence measurements in both cell types revealed that ROS levels remained unaltered in contact-free co-cultures (Fig. 4d), indicating that ROS transfer could only occur via a contact-dependent mechanism.

As it had been previously reported that haematopoietic stem cells can transfer ROS to stromal cells via gap junction to prevent senescence ^32^, we decided to examine gap junction genes in the transcriptome of MS-5 cultured alone and in co-culture. When individually examining the gap junction genes in MS-5 cells cultured alone vs co-culture, we found several gap junction genes upregulated in co-culture such as *Gja5, Gja8, Gjb5* and *Gjc2* (Fig. 4e). These results suggest that AML cells might establish gap junction interactions to transfer ROS to MS-5 cells when in co-culture. To test this hypothesis, we used the calcein-AM dye, which can only be transferred via gap junctions ^10^. We labelled stromal cells with calcein-AM, cultured them with unlabelled AML cells and analysed the fluorescence of AML cells after three hours. We found that, for the three AML cell lines tested, the percentage of cells that had incorporated the calcein-AM dye from MS-5 cells was larger than 80% (Fig. 4f, Supplementary Fig. 5c), indicating that AML cells can establish gap junctions with stromal cells when co-cultured in direct contact.

Next, we decided to confirm that AML cells can transfer ROS via gap junctions by inhibiting the gap junctions using carbenoxolone (CBX), a well-known gap junction inhibitor ^10, 33, 34^. We first confirmed that efficiency of inhibition by analysing the calcein-AM dye transfer in a control and treated co-culture of SKM-1 and MS-5 cells. The CBX treatment reduced the calcein-AM transfer more than 60% in co-culture (Fig. 4g). We then compared intracellular ROS levels in both AML and stromal cells treated with CBX and control. CBX treatment abrogated the decrease in ROS levels in AML cells (Fig. 4h) and the increase of ROS levels in stromal cells (Fig. 4i), indicating inhibition of ROS transfer in CBX treated co-cultures. Overall, this result provides strong evidence that ROS transfer occurs via gap junctions established between AML and stromal cells.

## Discussion

AML cells are known to interact and remodel niche cells through various mechanisms, including the secretion of soluble factors, cytokines or metabolites, resulting in a better support of AML cells at the expense of normal haematopoiesis. Yet, metabolic crosstalk between AML and stromal cells has not been reported before. Here, we have identified a novel metabolic communication between AML and stromal cells mediated by acetate. Our in vivo data shows high acetate levels in the bone marrow extracellular fluid of AML but not in healthy control mice, indicative of a potential role for acetate in AML development. Our in vitro data suggests that AML cells can modulate stromal cells into secreting acetate in co-culture, by rewiring stromal cell metabolism, and can then utilise the secreted acetate to feed their own TCA cycle to generate energy. Mechanistically, our data revealed that acetate secretion involves not only a higher glycolytic rate but also the non-enzymatic ROS-mediated conversion of pyruvate to acetate, as cells grown in thiamine-free media were still capable to produce acetate. Furthermore, our data indicates that AML cells can diminish their ROS levels by establishing gap junctions with stromal cells facilitating ROS transfer to stromal cells.

Studying the interactions between cancer cells and their microenvironment in terms of metabolism has become an exciting new field of cancer research. Our data indicates that AML cells can influence the metabolism of stromal cells causing increased acetate secretion, which was not observed in healthy counterparts. We presented several lines of evidence suggesting that stromal cells but not AML cells are responsible for acetate secretion: (i) stromal cells cultured alone secreted acetate, whilst AML cell lines and primary AML cells did not; (ii) after separating stromal cells from co-culture with AML cell lines, stromal cells continued to secrete acetate at a similar rate to that observed in co-culture; and, (iii) glycolysis was upregulated in stromal cells in co-culture, and glucose was found to be the precursor of the secreted acetate in co-culture.

Interestingly, acetate has been reported as an alternative fuel for cancer cells ^35, 36^, especially under low oxygen conditions or lipid depletion ^37–39^, but it has not previously been described to participate in crosstalk between any type of cancer cells and their microenvironment. Nonetheless, other monocarboxylate metabolites, such as lactate and alanine, have been reported to participate in different types of metabolic interplay between stromal and cancer cells ^40–42^. Our work reveals that aside from lactate and alanine, other monocarboxylate metabolites, such as acetate, can be utilised by leukaemic cells as a biofuel. Although acetate usage to feed the TCA cycle had already been described in AML ^43^ and other types of cancer ^38, 44^, this is, to our knowledge, the first report of AML cells utilising acetate secreted by stromal cells in co-culture.

We have also proposed a mechanism for the altered stromal cellular metabolism, involving increased glycolysis and the ROS-mediated chemical conversion of pyruvate to acetate. Transcriptomics showed that glycolysis is upregulated in stromal cells in co-culture, and tracer-based metabolism using [U-^13^C]glucose, demonstrated that more label gets incorporated in pyruvate and acetate in stromal cells in co-culture compared to either AML or stromal cells cultured alone. Enrichment of hypoxia genes and elevated Pdk expression have been reported to be related to higher glycolytic activity ^45–47^. Similarly, other cancer cells ^48–51^ have also been shown to modulate stromal metabolism by increasing glycolysis. Moreover, this mechanism was further supported by increasing and lowering ROS levels in co-culture. Lowering ROS levels recovered the acetate secretion phenotype of MS-5 cells cultured alone.

AML cells are known to exhibit high levels of ROS ^52, 53^. However, our data has shown that AML cells in co-culture present lower levels of ROS than AML cells cultured alone, which is in agreement with recent studies in primary AML and mesenchymal stromal cell co-cultures ^14, 24^. The current mechanisms to describe this phenomenon involve mitochondrial transfer and activation of glutathione-related antioxidant pathways ^14, 24^, although previous data on haematopoietic stem cells (HSCs) revealed that HSCs can directly transfer ROS via gap junctions to stromal cells ^32^. Interestingly, our data showed that the decrease in ROS levels was counteracted by inhibiting contact using a permeable membrane. We also found several gap junction genes upregulated in stromal cell types in co-culture. Moreover, we prevented ROS transfer from AML to stromal cells by using the gap junction inhibitor CBX ^10, 33, 34^. Thus, our results suggest that ROS transfer via gap junctions, at least partially, mediates the mechanism behind AML cells presenting lower levels of ROS in co-culture.

Overall, this work reveals a unique and novel metabolic communication between AML and stromal cells that involves acetate as the main crosstalk metabolite. We showed that AML cells are capable of modulating the metabolism of stromal cells by transferring ROS via gap junctions resulting in an increased secretion of acetate and its subsequent accumulation in the extracellular medium, which correlates with the observation of higher acetate levels in the bone marrow extracellular fluid of AML mice compared to control mice. Furthermore, we found that AML cells consume the secreted acetate and use it as an energy source, by fluxing it into the TCA cycle. We believe our findings provide a better understanding of how AML cells communicate with stromal cells and could serve as a basis for the development of novel therapeutic strategies to target AML cells by modulating gap junction formation as an adjuvant therapy.

## Materials and Methods

### Cell lines

The human AML cell lines (SKM-1, Kasumi-1 and HL-60), the mouse stromal cell line (MS-5) and the human cervical cancer cell line (HeLa) were cultured in RPMI 1640 media supplemented with 15% (v/v) FBS, 2 mmol/L L-glutamine and 100 U/ml Penicillin/Streptomycin (all from Thermo Fisher Scientific). Co-cultures were plated in a 4:1 AML-stromal ratio, and 750.000 cells/ml density of AML cells over confluent stromal cells. Prior to cell extraction for NMR, RNA collection or protein extraction, a suspension of 10 million AML cells was collected, the stromal layer was washed with PBS and was subject to mild trypsinisation with 1:5 dilution of 0.25% Trypsin 1 mM EDTA (Thermofisher) to remove attached residual leukaemic cells before completely detaching stromal cells with 0.25% Trypsin-EDTA.

### Primary patient samples

AML and PBMC primary specimens’ procedures were obtained in accordance with the Declaration of Helsinki at the University Medical Center Groningen, approved by the UMCG Medical Ethical Committee or at the University Hospital Birmingham NHS Foundation Trust, approved by the West Midlands – Solihull Research Ethics Committee (10/H1206//58).

Additional information about the primary AML samples used in this study can be found in Annex 1. Peripheral blood (PB) and bone marrow samples from AML patients and healthy donors were obtained in heparin-coated vacutainers. Mononuclear cells were isolated using Ficoll-Paque (GE Healthcare) and stored at −80°C.

For AML1-2 and PBMC1-2, samples were thawed and resuspended in newborn calf serum (NCS) supplemented with DNase I (20 Units/mL), 4 mM MgSO_4_ and heparin (5 Units/mL) and incubated on 37°C for 15 minutes (min). For AML1 and PBMC1, cells were sorted after thawing by fluorescence-activated cell sorting (FACS) for the CD34+CD38- population using 10 μL of anti-CD34 APC (560940, BD Biosciences), 10 μL of anti-CD38 FITC (555459, BD Biosciences) and 10 μL of DAPI (D1306, Thermo Fisher Scientific) per 10 million cells. Cells were sorted using a Sony SH800S (Sony) sorter. For AML2 and PBMC2, cells were thawed and the CD34+ population was sorted by magnetic-activated cell sorting (MACS) using 10 μL of anti-CD34 microbeads (130-046-702, Miltenyi) per million of expected CD34 cells following manufacturer’s protocol.

AML1-2, PBMC1-2 and MS-5 cells were cultured alone and in co-culture in a 4:1 AML/PBMC-stromal ratio and 500,000 cells/mL density in α-MEM (Gibco) with 12.5% (v/v) FCS (Sigma-Aldrich), 1% (v/v) Pen/Strep (Thermo Fisher Scientific), 12.5% (v/v) Horse serum (Invitrogen), 0.4% (v/v) β-mercaptoethanol (Merck Sharp & Dohme BV) and 0.1% hydrocortisone (H0888, Sigma-Aldrich). For AML1 and AML2 the medium was supplemented with 0.02 μg/mL of IL-3 (Sandoz), 0.02 μg/mL NPlate (Amgen) and 0.02 μg/mL of G-CSF (Amgen). For PBMC1 and PBMC2 the medium was supplemented with 100 ng/ml of human SCF (255-SC, Novus Biologicals), 100 ng/ml of NPlate, 100 ng/ml of FLT3 ligand (Amgen) and 20 ng/ml of IL-3. Samples of medium were collected at 0 and 48 hours.

AML3 - 7 and PBMC3 were thawed and kept in culture for 16-24 hours in Stem Span H3000 media (STEMCELL Technologies) supplemented with 50 μg/ml ascorbic acid (Sigma-Aldrich), 50 ng/ml human SCF (255-SC-010, R&D Systems), 10 ng/ml human IL-3 (203-IL-010, R&D Systems), 2 units/ml human-erythropoietin (100-64, PeproTech), 40 ng/ml insulin-like growth factor 1 (IGF-1) (100-12, PeproTech), 1 μM dexamethasone (D2915, Sigma-Aldrich) and 100 μg/ml primocin (ant-pm-2, InvivogenCD34+ cKit+ cells were purified using magnetic microbeads (130-046-702 (CD34) and 130-091-332 (CD117), Miltenyi Biotec). Cells were cultured in the supplemented Stem Span media for 24 hours prior to co-culture setting. Co-cultures were plated with a 4:1 leukaemic to stromal cells ratio and a 300,000 cells/ml density in supplemented Stem Span medium. Samples of leukaemic/healthy cells and MS-5 cells alone were also cultured in supplemented Stem Span media as controls. Samples of medium were collected at 0 and 48 hours.

### In vivo experiments

12 weeks old C57BL6/J female mice (The Jackson Laboratory) were lethally irradiated with 9Gy (2 doses of 4.5Gy separated by 3h) using an X-RAY source (Rad Source’s RS 2000). Mice were transplanted by intravenous injection 4h after with 2×10^6^ bone marrow (BM) nucleated cells isolated as previously described ^54^ from leukemic male mice heterozygous for MLL-AF9 knock-in fusion transgene or wild-type (WT) control male littermates ^55^. Male transgenic and WT control MLL-AF9 were purchased from The Jackson Laboratory (Stock No: 009079) and were 6 months old when euthanized to allow development of signs of leukemic transformation driven by the MLL-AF9 fusion oncogene in transgenic animals. MLL-AF9 expression in BM cells derived from transgenic donors results in development of AML with high blast counts in recipients. Recipients were euthanized 6 months after the transplant, bone marrow from femur and tibia was flushed with a syringe in 150 µL of cold PBS and centrifuged at 15000g for 10 minutes at 4°C. The supernatant made of bone marrow extracellular fluid (BMEF) was kept for analysis of acetate level. Experiments were conducted with the ethical approval of the Norwegian Food and Safety Authority. Animals were housed under specific opportunistic and pathogen free environment at the Animal Facility of the University of Oslo, Norway.

### Proliferation analysis using CFSE

The CellTrace™ carboxyfluorescein succinimidyl ester (CFSE) Cell Proliferation Kit (C34570, Invitrogen) was used to assess proliferation of AML cells following the manufacturer’s protocol. AML cells were stained and their fluorescence was assessed before dividing the bulk of cells into culturing them alone or with MS-5 cells for 48 hours. Small aliquots of cells after 24 and 48 hours were analysed by flow cytometry. Flow cytometry analysis was carried out in a CyAn ADP flow cytometer (Beckman Coulter). Data analysis was performed using the FlowJo software package (BD).

### Carbenoxolone treatment

Carbenoxolone (CBX) disodium salt (Sigma-Aldrich, C4790) was prepared fresh at 200 μM in cell culture medium. Cells were resuspended in the CBX medium and cultured alone or in co-culture for 24 hours prior to ROS or Calcein-AM dye transfer experiments.

### Cellular ROS measurements using DCFH-DA

Cellular ROS was measured by incubating cells with 100 µM 2′,7′-Dichlorofluorescin diacetate (DCFH-DA) (D6883, Sigma-Aldrich) in Hank’s Balanced Salt Solution (HBSS) (Thermo Fisher Scientific) at 37°C for 30 min protected from light. Cells were then harvested and stained with 5 μg/μL anti-human CD33 eFluor 450 (eBioscience, P67.6) for 30 min at 4°C before flow cytometry analysis as previously described.

### Thiamine-free medium comparison

Thiamine-free medium (R9011-01, United States Biological) was prepared as per manufacturer instructions and supplemented with 10% dialised FBS.

MS-5 cells were cultured in thiamine-free medium or control RPMI medium for 4 days prior to the experiment. Cells were then seeded with fresh thiamine-free or control RPMI medium and incubated for 24 hours. Samples of medium were collected at 0 and 24 hours and kept at −80°C.

### ROS-related treatments with H_2_O_2_ and NAC

SKM-1, Kasumi, HL-60 and MS-5 cells cultured alone and in co-culture were incubated for 24 hours in 50 μM H_2_O_2_ (Merck) complete cell culture medium, 5 mM N-acetylcysteine (NAC) (106425, Merck) complete cell culture medium or control medium. Samples of medium were collected at 0 and 24 hours and kept at −80°C.

### Calcein-AM dye transfer assay

Functional gap junction presence was evaluated using the fluorescent dye Calcein-AM green (Invitrogen, C1430) adapting a previously established protocol ^10^. MS-5 cells were stained with 500 nM Calcein-AM dye in complete cell culture medium for 1 hour at 37°C. Stained cells were washed with serum-free medium for 30 min at 37°C before being co-cultured with AML cells for 3h. AML cells were then harvested and stained with 5 μg/μL anti-human CD33 eFluor 450 (eBioscience, P67.6) for 30 min at 4°C before flow cytometry analysis as previously described. Calcein-AM dye transfer was quantified as the frequency of CD33^+^ and Calcein-AM^+^ cells.

### Tracer-based NMR experiments

[U-^13^C]Glucose (CC860P1, CortecNet) was added to RPMI 1640 medium without glucose (11879020, Merck) to a final concentration of 2 g/L (as in the complete cell culture medium) and was supplemented as usual with 15% (v/v) FBS, 2 mmol/L L-glutamine and 100 U/mL Pen/Strep. The medium was prepared fresh and was filtered with a 0.2 μm syringe filter (Sartorius) before each experiment.

4 mM sodium [2-^13^C]acetate (279315-1G, Sigma-Aldrich) was added to complete cell culture medium and the medium was filtered with a 0.2 μm syringe filter before each experiment. Cells were incubated for the time indicated in each experiment before separation of cells and/or metabolite extraction. **a**, **b**, **c**, **d**, **f**, **g**, **h** and **i** Unlabelled samples were prepared as a control for 2D-NMR experiments and to measure acetate consumption by adding unlabelled sodium acetate trihydrate (1.37012, Merck) to complete cell culture medium before metabolite extraction or collection of medium samples.

### Metabolite extraction

Suspension cells and detached adherent cells were washed with PBS before being rapidly resuspended in 400 µl of HPLC grade methanol pre-chilled on dry ice. Samples were transferred to glass vials and were subject to 10:8:10 methanol-water-chloroform extraction as described in (Saborano 2019). Polar phase samples were evaporated in a SpeedVac concentrator and were subsequently kept at −80°C.

### Sample preparation for NMR

Medium samples or bone marrow extracellular fluid samples were mixed with a D_2_O phosphate buffer containing TMSP ((3-trimethylsilyl)propionic-(2,2,3,3-d_4_)-acid sodium salt) and NaN_3_ to a final concentration of 0.1 M phosphate and 0.5 mM TMSP ((3-trimethylsilyl)propionic-(2,2,3,3-d_4_)-acid sodium salt) and 1.5 mM NaN_3_ (all from Sigma) and transferred to 3 mm NMR tubes (Cortecnet).

Dried polar extracts for tracer-based NMR experiments were reconstituted in 50 μL of 0.1 M phosphate buffer in 100% D_2_O with 3 mM NaN_3_ and 0.5 mM TMSP. Samples were sonicated for 10 min and transferred to 1.7 mm NMR tubes using a micro pipet system. Samples were prepared freshly before the acquisition.

### NMR acquisition and analysis

All NMR data were acquired on Bruker 600MHz spectrometers equipped with Avance-III consoles using cooled Bruker SampleJet autosamplers. For media samples, a 5mm triple resonance cryoprobe (TCI) z-axis pulsed field gradient (PFG) cryogenic probe was used, and for cell extracts, a TCI 1.7mm z-PFG cryogenic probe was used. Probes were equipped with a cooled SampleJet autosampler (Bruker) and automated tuning and matching.

For medium samples, spectra were acquired at 300K using 1D ^1^H-NOESY (Nuclear Overhauser effect spectroscopy) pulse sequence with pre-saturation water suppression (noesygppr1d, standard pulse sequence from Bruker). The spectral width was 12 ppm, the number of data points was TD 32,768, the interscan delay was 4 sec and the NOE mixing time was 10 msec. The ^1^H carrier was set on the water frequency and the ^1^H 90° pulse was calibrated at a power of 0.256 W and had a typical length of ca 7-8 μs. 64 scans and 8 dummy scans were acquired and the total experimental time was 7.5 min. Spectra were processed using the MetaboLab ^56^ software within the MATLAB environment (MathWorks). A 0.3Hz exponential apodization function was applied to FIDs followed by zero-filling to 131,072 data points prior to Fourier transformation. The chemical shift was calibrated by referencing the TMSP signal to 0 ppm and spectra were manually phase corrected. The baseline was corrected by applying a spline baseline correction, the water and edge regions of the spectra were excluded before scaling the spectra using a probabilistic quotient normalization (PQN). Chenomx 7.0 software (Chenomx Inc.) and the human metabolome database (HMDB) were used to assign the metabolites present in the acquired spectra. Metabolite signal intensities were obtained directly from the spectra and were normalised to a control medium sample obtained at time 0 hours. For bone marrow extracellular fluid samples, metabolite signal areas were obtained directly from the spectra.

For ^13^C-filtered ^1^H-NMR experiments, [U-^13^C]glucose labelled medium samples were analysed with ^13^C-filtered ^1^H-NMR spectroscopy as described in ^57^. Spectra were acquired at 300 K using a double gradient BIRD filter pulse sequence developed in-house ^57^. A pulse program combining the ^1^H[^12^C] and the all-^1^H experiments in scan-interleaved mode was used. The difference between the two FIDs gave the ^1^H[^13^C] signal. The spectral width was 12 ppm, the number of data points was 16,384 in each dimension and the relaxation delay was 5.3 sec. 256 scans with 64 dummy scans were acquired and the experimental time was 15 min. ^13^C-filtered ^1^H-NMR spectra were processed in Topspin 4.0.5 (Bruker). Spectra were zero filled to 32,768 data points before Fourier transformation. Phase correction was applied to the ^1^H[-^12^C] and the all-^1^H spectra and the difference ^1^H[^13^C] spectrum was obtained by aligning on chemical shift. Metabolites were selected in the ^1^H[^13^C] spectrum and integrated in all the spectra (^1^H[^12^C], ^1^H[^13^C] and all-^1^H). Label incorporations (^13^C percentages) were calculated by dividing the signal areas obtained in the ^1^H[^13^C] spectrum by the ones obtained in the all-^1^H spectrum.

For ^1^H-^13^C-HSQC experiments, spectra were acquired using a modified version of Bruker’s hsqcgphprsp pulse program, with additional gradient pulses during the INEPT echo periods and using soft 180° pulses for ^13^C. For the ^1^H dimension, the spectral width was 13.018 ppm with 2048 complex points. For the ^13^C dimension, the spectral width was 160 ppm with 2048 complex points. 2 scans were acquired per spectrum and with an interscan delay of 1.5 sec. Non-uniform sampling (NUS) with a 25% sampling schedule (generated using the Wagner’s schedule generator (Gerhard Wagner Lab, Harvard Medical School) with a tolerance of 0.01 and default values for the other parameters) was used with 4096 increments yielding 8192 complex points after processing. The total experimental time was 4 hours. Spectra were processed and phased with NMRPipe (National Institute of Standards and Technology of the U.S.) (Appendix 8.1). MetaboLab was used to reference the chemical shift using the signal for the methyl group of L-lactic acid, at 1.31/22.9 ppm in the ^1^H and ^13^C dimensions, respectively. Identification of metabolites in 2D spectra was carried out using the MetaboLab software which includes a chemical shift library for ca. 200 metabolites. Intensities were obtained from signals in the spectra and were corrected for differences in cell numbers contributing to the sample as follows: A ^1^H-NMR spectrum was acquired for each sample and the total metabolite area of this spectrum after removal of the water and TMSP reference signal was calculated in MetaboLab. The intensities in the 2D spectra were then divided by the total metabolite area of the corresponding ^1^H-NMR spectrum. To obtain the % of ^13^C in a metabolite, the normalized intensity of a certain carbon in the labelled sample was divided by the normalized intensity of the same carbon in the unlabelled sample and was multiplied by the natural abundance of ^13^C (1.1%).

### RNA extraction and sequencing

MS-5 cells were cultured alone or in co-culture with SKM-1 cells for 24 hours. Cells were separated and washed with PBS prior to RNA extraction with TRIzol (Gibco) according to the manufacturer’s protocol. RNA was purified with RNeasy Plus Micro kit. Samples were sent to Theragenetex to be sequenced with Novaseq 150bp PE with 40M reads.

### Real-time PCR

Samples of RNA from SKM-1 and MS-5 co-cultures were collected and extracted using Trizol (Invitrogen) following manufacturer’s protocol. cDNA was synthesized using the M-MLV reverse transcriptase (Promega) according to manufacturer’s instructions. For gene expression analysis, qRT-PCR of *Hk2* (NM_013820), *Pdhx* (NM_175094), *Pdk1* (NM_172665) and *Pdk2* (NM_133667) (all KiCqStart™ primers KSPQ12012, Sigma Aldrich) were carried out using the SYBRGreen Master mix (Thermo Fisher Scientific) and qRT-PCR of *B2m* (NM_009735.3, TaqMan® assays, Thermo Fisher Scientific) was performed using TaqMan PCR Master Mix (Thermo Fisher Scientific). Reactions were performed in a Stratagene Mx3000P and were run in triplicate. Relative gene expression was calculated following the 2^−ΔΔCt^ method relative to the expression of *B2m*.

### RNA sequencing

RNA samples of MS-5 cells cultured alone and in co-culture with SKM-1 cells extracted using Trizol were purified using the Rneasy Plus Micro kit (Qiagen) according to manufacturer’s protocol. Transcriptome analysis was performed by Theragen Etex Co., LTD. (www.theragenetex.net). cDNA libraries were prepared with the TruSeq Stranded mRNA Sample Prep Kit (Illumina) and RNA sequencing was performed in a HiSeq2500 platform (Illumina). Quality control metrics were obtained with FastQC 0.11.7 software (Babraham Bioinformatics). To quantify transcript abundances using the Kallisto 0.43.0 software (Patcher Lab), read counts were aligned to the GRCm38 mouse reference genome cDNA index (Ensembl rel.67). Gene-level differential expression analysis was carried out with the R statistical package Sleuth 0.30.0 comparing the expression of cells cultured alone vs co-culture. Differentially expressed genes were calculated using the Wald statistical test, correcting for multiple testing with the Benjamin-Hochberg method. The false discovery rate (FDR) threshold was set at 1% (q-values < 0.01). Ensembl gene transcripts were annotated to Entrez IDs and official gene symbols with the R statistical package BioMart 2.40.3. To normalise for sequencing depth and gene length, transcripts per million (TPM) expression values were calculated. Gene Set Enrichment Analysis (GSEA) was performed with the R statistical package fgsea 1.10.0. The collection of hallmark gene sets from the Molecular Signature Database was used for the GSEA, setting the FDR threshold at 5%. Data was deposited in GEO (GSE163478).

## Acknowledgements

N. Vilaplana-Lopera, G. Papatzikas, A. Cunningham, and A. Erdem were supported by the EU grant HaemMetabolome H2020-MSCA-ITN-2015-675790. U. Günther, P. Garcia, J. J. Schuringa, J.-B. Cazier, and F. Schnütgen acknowledge support from the European Commission (HaemMetabolome [EC-675790]). This work was further supported by the Deutsche Forschungsgemeinschaft (DFG, German Research foundation) SFB815, TP A10 (F.S.). We also acknowledge the Wellcome Trust for supporting access to NMR instruments at the Henry Wellcome Building for Biomolecular NMR in Birmingham (grant number 208400/Z/17/Z). The mouse work was supported by a joint meeting grant of the Northern Norway Regional Health Authority and UiT (2014/5668) to L. Arranz. We thank A. Villatoro and L.M. Gonzalez for technical assistance.

## Author Contributions

N.V-L. conceived and performed the experiments, acquired, analysed and interpreted the data, and wrote the manuscript. V.C., R.A., E.G., A.C., A.E., F.S., M.R., L.A. performed the experiments and edited the manuscript. G.P. analysed the RNA-seq data. J.B helped with the experimental design. S.P. and M.R. provided with patient samples. V.C. and L.A. provided mouse bone marrow extracellular fluid samples. J.J.S. provided patient samples, helped with the experimental design, interpretation and critical discussion of the data and edited the manuscript. U.G and P.G. conceived and performed the experiments, acquired and interpreted the data, wrote the manuscript, and managed the project.

## EXTENDED DATA

**Extended Data Figure 1.**
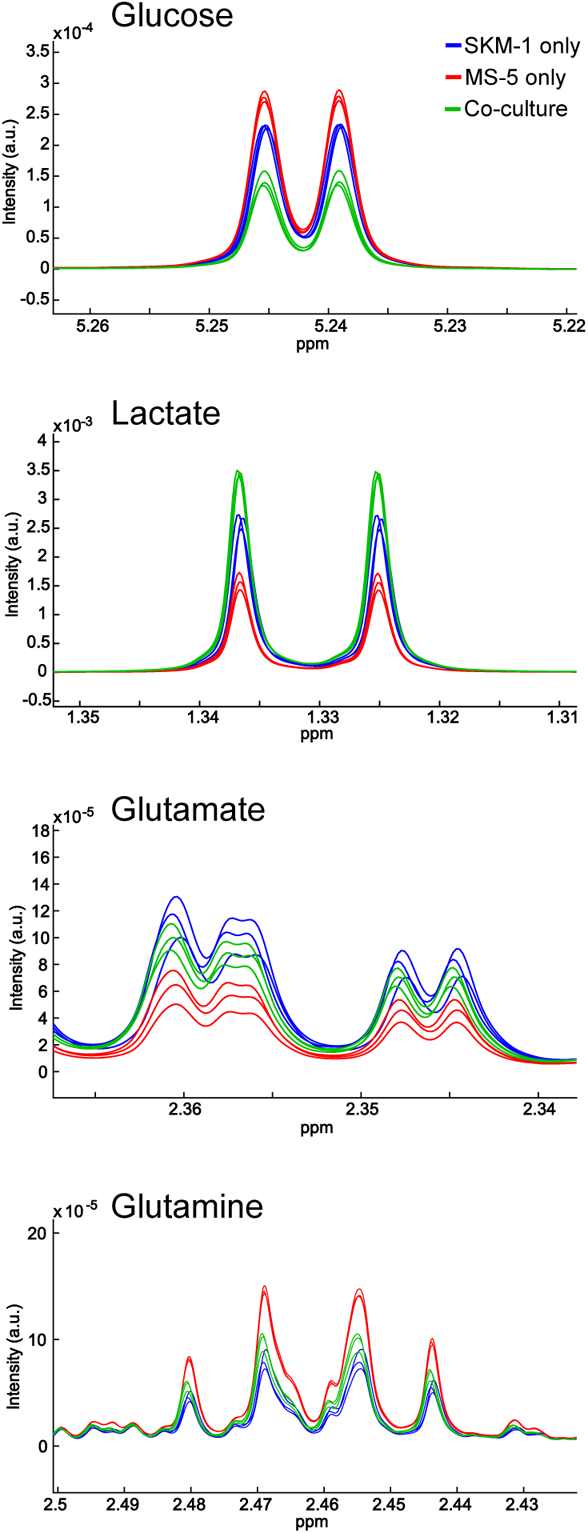
Other extracellular metabolite levels in co-cultures with SKM-1 and MS-5 cells. Sections of ^1^H-NMR spectra from extracellular medium samples of SKM-1 cells cultured alone (blue), MS-5 cells cultured alone (red) and SKM-1 and MS-5 cells in co-culture (green) after 24 hours of culture, corresponding to glucose, lactate, glutamate and glutamine.

**Extended Data Figure 2.**
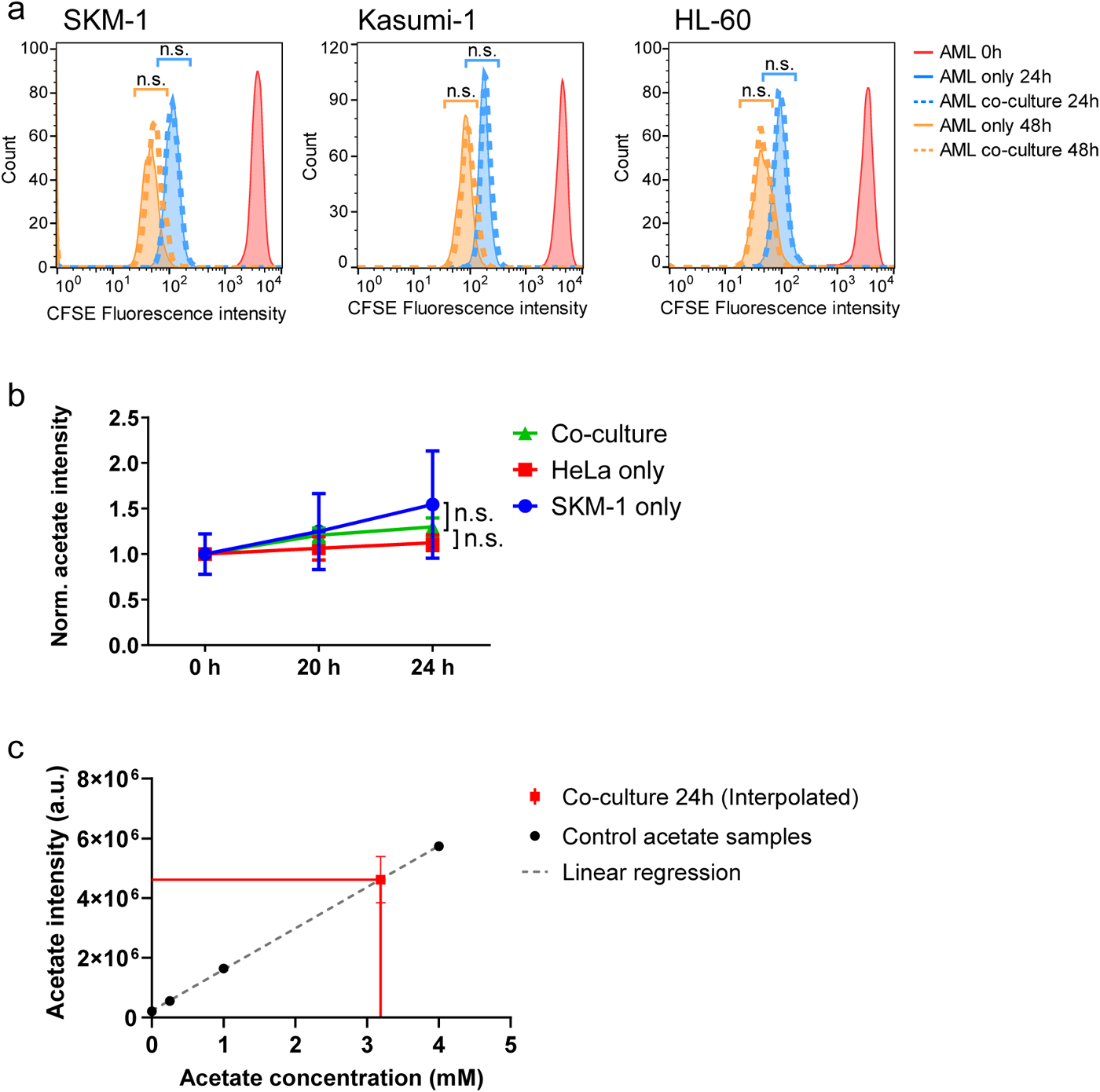
Proliferation in co-culture, co-culturing AML cells with an unrelated cell line (HeLa) and titration of acetate concentration in co-culture. (**a**) CFSE cell proliferation analysis in AML cell lines alone or in co-culture with MS-5 cells. The population of living cells was gated, and 1500 cells for each condition were randomly selected and plotted. The geometric mean for each population and time point was compared between cells alone or in co-culture by performing an unpaired unpaired Student’s t-test and p-values were represented as n.s. for not significant. Each histogram is representative of n=3 independent experiments. (**b**) Extracellular acetate levels in SKM-1 cells cultured alone (blue), HeLa cells cultured alone (red) and AML and HeLa cells in co-culture (green) at 0, 20 and 24 hours. Each point represents the mean of n=3 independent experiments and error bars represent standard deviation. An unpaired Student’s t-test was applied for each condition (black brackets) and p-values are represented by n.s. for not significant. (**c**) Linear regression of acetate concentrations and detected intensities in ^1^H-NMR spectra (Intensity=1.38·10^6^ ± 1.6·10^4^ x Acetate concentration + 2.27·10^5^ ± 3.3·10^4^; R^2^=0.9987). The intensities detected in samples of co-culture were interpolated to obtain the estimate acetate concentration in co-culture (3.09 mM). Each point represents the mean of n=3 independent experiments and error bars represent standard deviation.

**Extended Data Figure 3.**
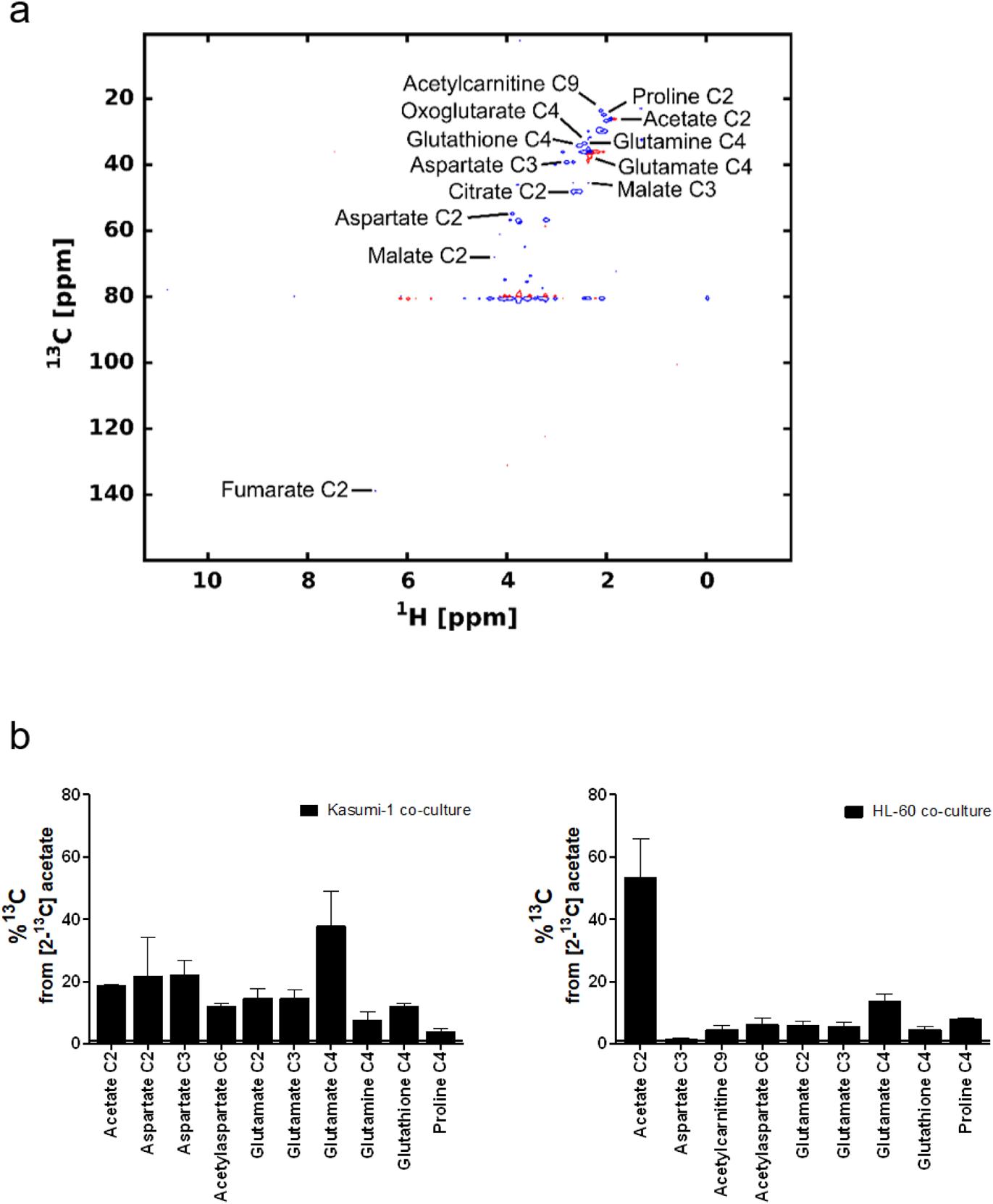
Label incorporation from [2-^13^C]acetate in SKM-1 and MS-5 cells cultured alone. (**a**) Example of metabolites assigned in a ^1^H-^13^C-HSQC spectrum from a polar extract from SKM-1 cells cultured with [2-^13^C]acetate for 2 hours. ^13^C percentages were calculated in 13 carbons from 11 different metabolites. The exact chemical shifts for each carbon are: acetate C2, 1.91 ppm (^1^H) – 26.1 ppm (^13^C); aspartate C2, 3.89 ppm (^1^H) – 55.0 ppm (^13^C); aspartate C3, 2.82 ppm (^1^H) – 39.4 ppm (^13^C); acetylcarnitine C9, 2.15 ppm (^1^H) – 23.4 ppm (^13^C); citrate C2, 2.52 ppm (^1^H) – 48.2 ppm (^13^C); fumarate C2, 6.51 ppm (^1^H) – 138.2 ppm (^13^C); glutamate C4, 2.34 ppm (^1^H) – 36.1 ppm (^13^C); glutamine C4, 2.44 ppm (^1^H) –33.7 ppm (^13^C); oxoglutarate C4, 2.43 ppm (^1^H) – 33.4 ppm (^13^C); glutathione C4, 2.55 ppm (^1^H) – 34.2 ppm (^13^C); malate C2, 4.28 ppm (^1^H) – 73.2 ppm (^13^C); malate C3, 2.68 ppm (^1^H) – 45.4 ppm (^13^C); and proline C4, 2 ppm (^1^H) – 26.6 ppm (^13^C). B) ^13^C percentages on polar metabolites in Kasumi-1 and HL-60 cells in co-culture with MS-5 cells. Cells were co-cultured for 24 hours before the addition of extra 4 mM sodium [2-^13^C]acetate. Bars represent the mean of the ^13^C percentage and error bars represent the standard deviation of n=3 independent experiments. ^13^C natural abundance is represented as a black bar at %^13^C = 1.1.

**Extended Data Figure 4.**
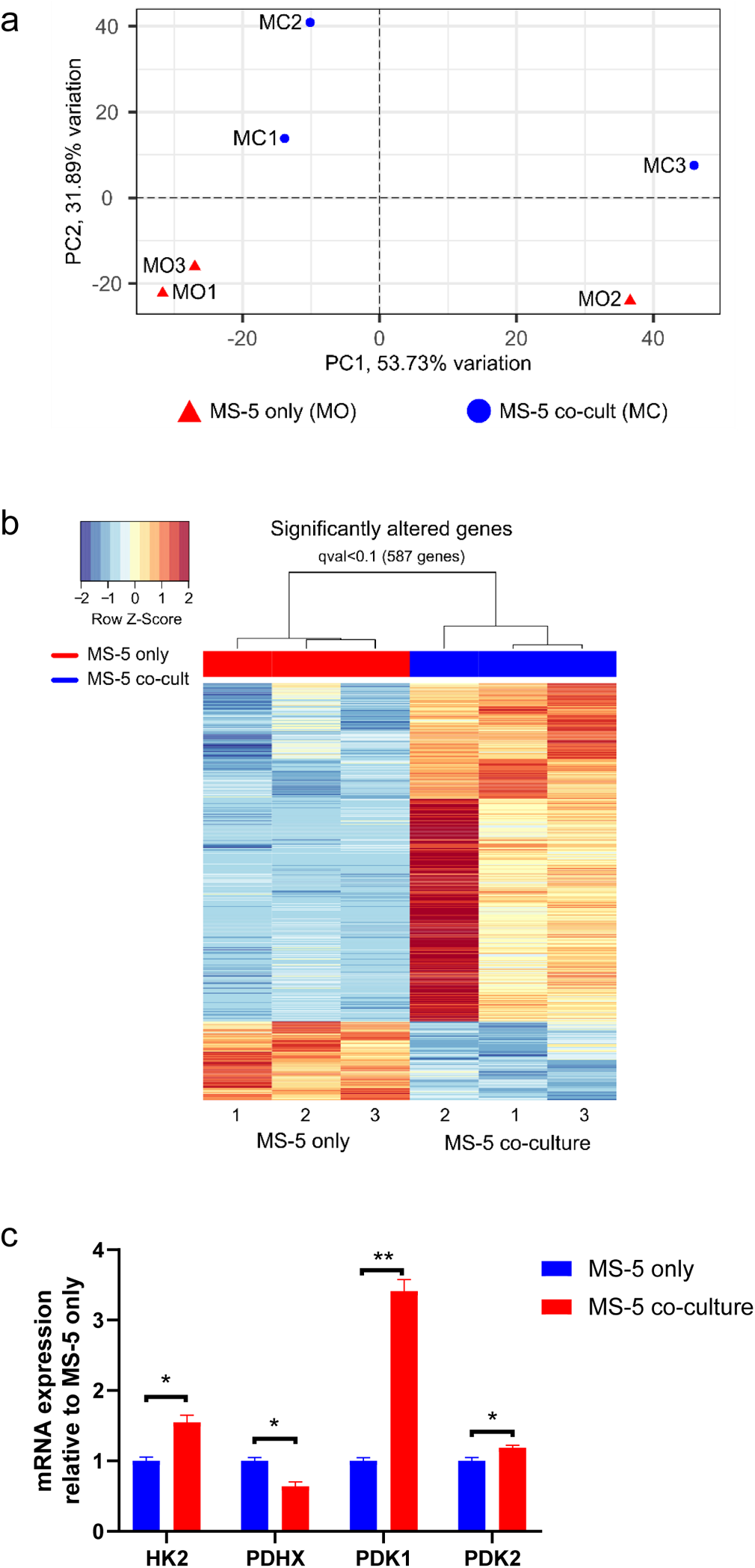
PCA component analysis, heat map of differentially expressed genes and qPCR for MS-5 cells cultured alone and in co-culture with SKM-1 cells. (**a**) PCA plot showing the clustering of individual samples of MS-5 cells cultured alone (MO) and MS-5 cells in co-culture (MC). The x and y axis values represent the variation between the sample groups. Generated using Sleuth 0.30.0 R statistical package. (**b**) Heat map of differentially expressed genes in MS-5 only (red) vs MS-5 co-culture (blue) calculated using the Wald statistical test, correcting for multiple testing comparison employing the Benjamini-Hochberg method using a false discovery rate threshold of 1% (q-value<0.01). Generated using Sleuth 0.30.0 R statistical package. (**c**) Quantitative PCR mRNA expression values for RNA sequencing validation of MS-5 cells cultured alone and in co-culture with SKM-1 cells. The gene set chosen was *Hk2*, *Pdhx*, *Pdk1* and *Pdk2*. mRNA quantification was normalized to *B2m* house-keeping gene. Bars represent the mean and error bars represent the SEM of three independent experiments. An unpaired Student’s t-test was applied for each gene and p-values are represented by * for p-value<0.05, ** for p-value<0.01 and *** for p-value<0.001.

**Extended Data Figure 5.**
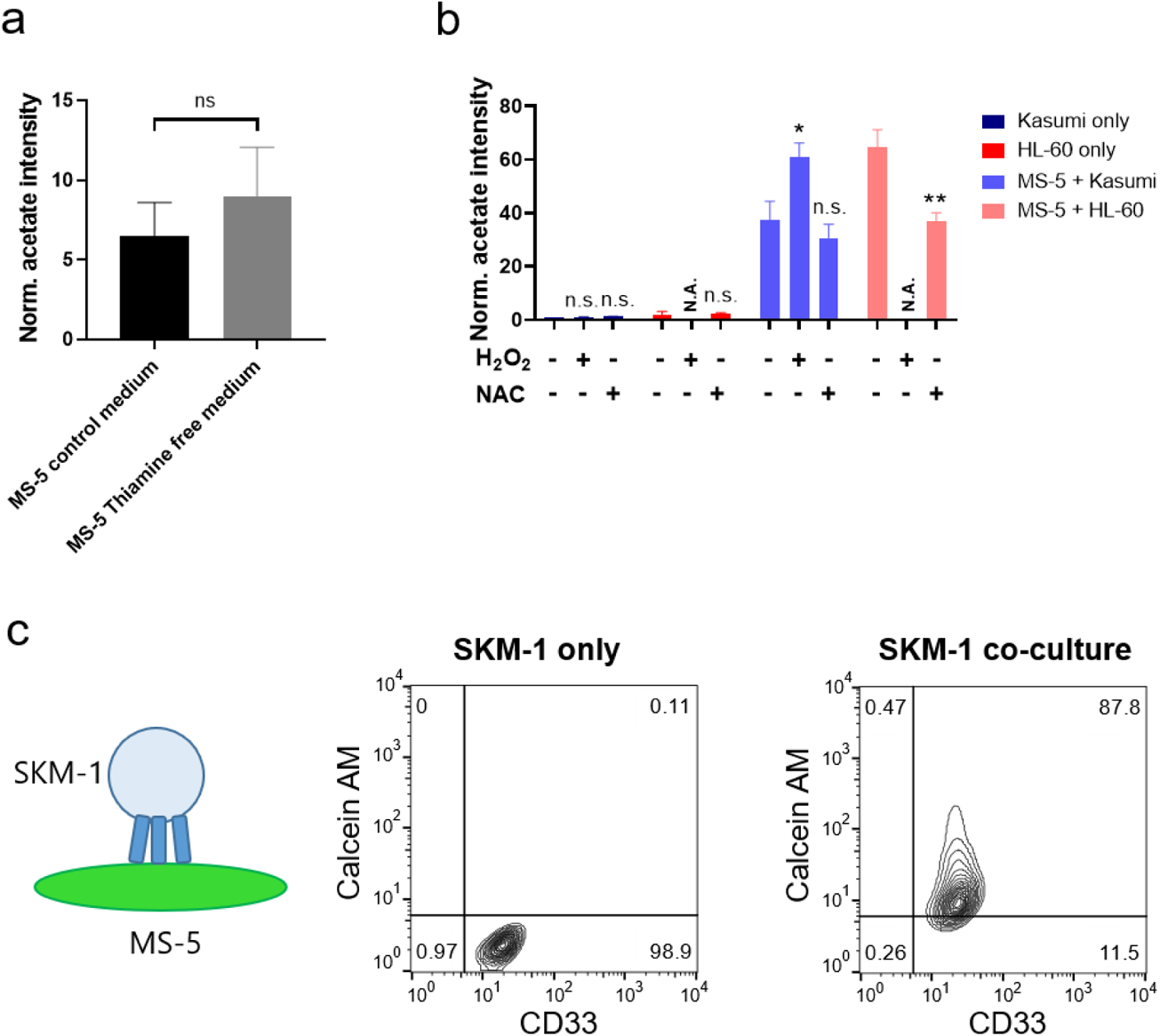
Acetate secretion in thiamine free medium and after modulating ROS levels and calcein-AM transfer through GAP junctions in AML-stromal co-cultures. (**a**) Extracellular acetate levels in MS-5 cells cultured in control medium (black) or in thiamine-depleted medium (grey) after 24 hours. (**b**) Extracellular acetate levels in Kasumi (navy blue) and HL-60 (red) cultured alone and in co-culture (light blue and light red) for 24 hours in a control medium, medium with 50 µM H_2_O_2_ or medium with 5 mM NAC. Acetate levels for HL-60 cells cultured alone or under co-culture with 50 µM H_2_O_2_ are not shown as treatment 50 µM H_2_O_2_ severely impaired their viability. N.A. = not analysed. For A and B each point represents the mean of n=3 independent experiments and error bars represent standard deviation. An unpaired Student’s t-test was applied for each condition (black brackets) and p-values are represented by n.s. for not significant. * for p-value<0.05 and ** for p-value<0.01. (**c**) Representative flow cytometry diagrams showing CD33 and calcein-AM green fluorescence in SKM-1 cells cultured alone or SKM-1 cells in co-culture with Calcein-AM-stained MS-5 cells after 3 hours. Frequencies of gated cell populations are indicated.

**Extended Data Table 1.** Primary AML samples’ additional information

| Patient | Tissue of origin | Type of AML (FAB) | Karyotype | Other mutations | Risk |
| --- | --- | --- | --- | --- | --- |
| AML1 | PB | M5 | Normal | FLT3-ITD; NPM1;<br>DNMTA3; IDH2 | Adv |
| AML2 | PB | M1 | Unknown | - | Adv |

**Samples obtained from University Hospital Birmingham NHS Foundation Trust, UoB** (used in figure 1e and 4c)
| Patient | Tissue of origin | Type of disease | Karyotype | Other mutations |
| --- | --- | --- | --- | --- |
| AML3 | PB | Secondary AML from MDS | 46,XY | TET2; SRSF2;<br>RUNX1; CEBPA;<br>ASXL1 |
| AML4 | PB | Secondary AML from MDS | 46,XY | NPM1 neg<br>FLT3 neg |
| AML5 | PB | AML M1 (FAB) | 46 XX | FLT3-ITD, 2xWT1,<br>CEBPA |
| AML6 | PB | AML M2 (FAB) | 46 XX | - |
| AML7 | PB | AML M4 (FAB) | 46 XY | - |

## References

1. Kreitz, J., et al. Metabolic Plasticity of Acute Myeloid Leukemia. Cells 8 (2019).

2. Schelker, R.C., et al. TGF-β1 and CXCL12 modulate proliferation and chemotherapy sensitivity of acute myeloid leukemia cells co-cultured with multipotent mesenchymal stromal cells. Hematology 23, 337–345 (2018).

3. Passaro, D., et al. Increased Vascular Permeability in the Bone Marrow Microenvironment Contributes to Disease Progression and Drug Response in Acute Myeloid Leukemia. Cancer Cell 32, 324–341 e326 (2017).

4. Zeng, Z., et al. Inhibition of CXCR4 with the novel RCP168 peptide overcomes stroma-mediated chemoresistance in chronic and acute leukemias. Mol Cancer Ther 5, 3113–3121 (2006).

5. Carey, A., et al. Identification of Interleukin-1 by Functional Screening as a Key Mediator of Cellular Expansion and Disease Progression in Acute Myeloid Leukemia. Cell reports 18, 3204–3218 (2017).

6. Zhang, T.Y., et al. IL-6 blockade reverses bone marrow failure induced by human acute myeloid leukemia. 12, eaax5104 (2020).

7. Wang, B., et al. Exosomes derived from acute myeloid leukemia cells promote chemoresistance by enhancing glycolysis-mediated vascular remodeling. J Cell Physiol 234, 10602–10614 (2019).

8. Kumar, B., et al. Acute myeloid leukemia transforms the bone marrow niche into a leukemia-permissive microenvironment through exosome secretion. Leukemia 32, 575–587 (2018).

9. Hornick, N.I., et al. AML suppresses hematopoiesis by releasing exosomes that contain microRNAs targeting c-MYB. Science signaling 9, ra88 (2016).

10. Kouzi, F., et al. Disruption of gap junctions attenuates acute myeloid leukemia chemoresistance induced by bone marrow mesenchymal stromal cells. Oncogene 39, 1198–1212 (2020).

11. Omsland, M., Bruserud, Ø., Gjertsen, B.T. & Andresen, V. Tunneling nanotube (TNT) formation is downregulated by cytarabine and NF-κB inhibition in acute myeloid leukemia (AML). Oncotarget 8, 7946–7963 (2017).

12. Ye, H., et al. Leukemic Stem Cells Evade Chemotherapy by Metabolic Adaptation to an Adipose Tissue Niche. Cell Stem Cell 19, 23–37 (2016).

13. Moschoi, R., et al. Protective mitochondrial transfer from bone marrow stromal cells to acute myeloid leukemic cells during chemotherapy. Blood 128, 253–264 (2016).

14. Forte, D., et al. Bone Marrow Mesenchymal Stem Cells Support Acute Myeloid Leukemia Bioenergetics and Enhance Antioxidant Defense and Escape from Chemotherapy. Cell Metab 32, 829–843 e829 (2020).

15. Shafat, M.S., et al. Leukemic blasts program bone marrow adipocytes to generate a protumoral microenvironment. Blood 129, 1320–1332 (2017).

16. Tabe, Y., et al. Bone Marrow Adipocytes Facilitate Fatty Acid Oxidation Activating AMPK and a Transcriptional Network Supporting Survival of Acute Monocytic Leukemia Cells. Cancer Res 77, 1453–1464 (2017).

17. Baccelli, I., et al. Mubritinib Targets the Electron Transport Chain Complex I and Reveals the Landscape of OXPHOS Dependency in Acute Myeloid Leukemia. Cancer Cell 36, 84–99.e88 (2019).

18. Pollyea, D.A., et al. Venetoclax with azacitidine disrupts energy metabolism and targets leukemia stem cells in patients with acute myeloid leukemia. Nat Med 24, 1859–1866 (2018).

19. Molina, J.R., et al. An inhibitor of oxidative phosphorylation exploits cancer vulnerability. Nature Medicine 24, 1036–1046 (2018).

20. Lagadinou, E.D., et al. BCL-2 inhibition targets oxidative phosphorylation and selectively eradicates quiescent human leukemia stem cells. Cell Stem Cell 12, 329–341 (2013).

21. Farge, T., et al. Chemotherapy-Resistant Human Acute Myeloid Leukemia Cells Are Not Enriched for Leukemic Stem Cells but Require Oxidative Metabolism. Cancer Discov 7, 716–735 (2017).

22. Li, L., et al. Altered Hematopoietic Cell Gene Expression Precedes Development of Therapy-Related Myelodysplasia/Acute Myeloid Leukemia and Identifies Patients at Risk. Cancer Cell 20, 591–605 (2011).

23. Hole, P.S., et al. Overproduction of NOX-derived ROS in AML promotes proliferation and is associated with defective oxidative stress signaling. Blood 122, 3322–3330 (2013).

24. Vignon, C., et al. Involvement of GPx-3 in the Reciprocal Control of Redox Metabolism in the Leukemic Niche. Int J Mol Sci 21 (2020).

25. Itoh, K., et al. Reproducible establishment of hemopoietic supportive stromal cell lines from murine bone marrow. Exp Hematol 17, 145–153 (1989).

26. Childress, C.C., Sacktor, B. & Traynor, D.R. Function of carnitine in the fatty acid oxidase-deficient insect flight muscle. J Biol Chem 242, 754–760 (1967).

27. Stephens, F.B., Constantin-Teodosiu, D. & Greenhaff, P.L. New insights concerning the role of carnitine in the regulation of fuel metabolism in skeletal muscle. J Physiol 581, 431–444 (2007).

28. Liu, X., et al. Acetate Production from Glucose and Coupling to Mitochondrial Metabolism in Mammals. Cell 175, 502–513 e513 (2018).

29. Vysochan, A., Sengupta, A., Weljie, A.M., Alwine, J.C. & Yu, Y. ACSS2-mediated acetyl-CoA synthesis from acetate is necessary for human cytomegalovirus infection. Proc Natl Acad Sci U S A 114, E1528–E1535 (2017).

30. Tiziani, S., et al. Metabolomic profiling of drug responses in acute myeloid leukaemia cell lines. PLoS One 4, e4251 (2009).

31. Nogueira-Pedro, A., et al. Hydrogen peroxide (H2O2) induces leukemic but not normal hematopoietic cell death in a dose-dependent manner. Cancer Cell International 13, 123 (2013).

32. Taniguchi Ishikawa, E., et al. Connexin-43 prevents hematopoietic stem cell senescence through transfer of reactive oxygen species to bone marrow stromal cells. Proc Natl Acad Sci U S A 109, 9071–9076 (2012).

33. Davidson, J.S., Baumgarten, I.M. & Harley, E.H. Reversible inhibition of intercellular junctional communication by glycyrrhetinic acid. Biochem Biophys Res Commun 134, 29–36 (1986).

34. Davidson, J.S. & Baumgarten, I.M. Glycyrrhetinic acid derivatives: a novel class of inhibitors of gap-junctional intercellular communication. Structure-activity relationships. J Pharmacol Exp Ther 246, 1104–1107 (1988).

35. Lyssiotis, C.A. & Cantley, L.C. Acetate fuels the cancer engine. Cell 159, 1492–1494 (2014).

36. Comerford, S.A., et al. Acetate dependence of tumors. Cell 159, 1591–1602 (2014).

37. Gao, X., et al. Acetate functions as an epigenetic metabolite to promote lipid synthesis under hypoxia. Nat Commun 7, 11960 (2016).

38. Schug, Z.T., et al. Acetyl-CoA synthetase 2 promotes acetate utilization and maintains cancer cell growth under metabolic stress. Cancer Cell 27, 57–71 (2015).

39. Yoshii, Y., et al. Cytosolic acetyl-CoA synthetase affected tumor cell survival under hypoxia: the possible function in tumor acetyl-CoA/acetate metabolism. Cancer Sci 100, 821–827 (2009).

40. Sousa, C.M., et al. Pancreatic stellate cells support tumour metabolism through autophagic alanine secretion. Nature 536, 479–483 (2016).

41. Sonveaux, P., et al. Targeting lactate-fueled respiration selectively kills hypoxic tumor cells in mice. J Clin Invest 118, 3930–3942 (2008).

42. Whitaker-Menezes, D., et al. Evidence for a stromal-epithelial “lactate shuttle” in human tumors: MCT4 is a marker of oxidative stress in cancer-associated fibroblasts. Cell Cycle 10, 1772–1783 (2011).

43. Saborano, R., et al. A framework for tracer-based metabolism in mammalian cells by NMR. Sci Rep 9, 2520 (2019).

44. Mashimo, T., et al. Acetate is a bioenergetic substrate for human glioblastoma and brain metastases. Cell 159, 1603–1614 (2014).

45. Testa, U., Labbaye, C., Castelli, G. & Pelosi, E. Oxidative stress and hypoxia in normal and leukemic stem cells. Experimental Hematology 44, 540–560 (2016).

46. Kocabas, F., et al. Hypoxic metabolism in human hematopoietic stem cells. Cell & bioscience 5, 39 (2015).

47. Takubo, K., et al. Regulation of glycolysis by Pdk functions as a metabolic checkpoint for cell cycle quiescence in hematopoietic stem cells. Cell Stem Cell 12, 49–61 (2013).

48. Shan, T., et al. Cancer-associated fibroblasts enhance pancreatic cancer cell invasion by remodeling the metabolic conversion mechanism. Oncol Rep 37, 1971–1979 (2017).

49. Migneco, G., et al. Glycolytic cancer associated fibroblasts promote breast cancer tumor growth, without a measurable increase in angiogenesis: evidence for stromal-epithelial metabolic coupling. Cell Cycle 9, 2412–2422 (2010).

50. Pavlides, S., et al. The reverse Warburg effect: aerobic glycolysis in cancer associated fibroblasts and the tumor stroma. Cell Cycle 8, 3984–4001 (2009).

51. Cruz-Bermudez, A., et al. Cancer-associated fibroblasts modify lung cancer metabolism involving ROS and TGF-beta signaling. Free Radic Biol Med 130, 163–173 (2019).

52. Hole, P.S., et al. Overproduction of NOX-derived ROS in AML promotes proliferation and is associated with defective oxidative stress signaling. Blood 122, 3322–3330 (2013).

53. Robinson, A.J., et al. Reactive Oxygen Species Drive Proliferation in Acute Myeloid Leukemia via the Glycolytic Regulator PFKFB3. Cancer Res 80, 937–949 (2020).

54. Arranz, L., et al. Neuropathy of haematopoietic stem cell niche is essential for myeloproliferative neoplasms. Nature 512, 78–81 (2014).

55. Corral, J., et al. An Mll-AF9 fusion gene made by homologous recombination causes acute leukemia in chimeric mice: a method to create fusion oncogenes. Cell 85, 853–861 (1996).

56. Ludwig, C. & Gunther, U.L. MetaboLab--advanced NMR data processing and analysis for metabolomics. BMC Bioinformatics 12, 366 (2011).

57. Reed, M.A.C., Roberts, J., Gierth, P., Kupce, E. & Gunther, U.L. Quantitative Isotopomer Rates in Real-Time Metabolism of Cells Determined by NMR Methods. Chembiochem 20, 2207–2211 (2019).

